# A quantitative characterization of early neuron generation in the developing zebrafish telencephalon

**DOI:** 10.1101/2023.04.18.537159

**Authors:** Glòria Casas Gimeno, Ekaterina Dvorianinova, Carla-Sophie Lembke, Emma SC Dijkstra, Hussam Abbas, Yuanyuan Liu, Judith TML Paridaen

## Abstract

The adult brain is made up of anatomically and functionally distinct regions with specific neuronal compositions. At the root of this neuronal diversity are neural stem and progenitor cells (NPCs) that produce many neurons throughout embryonic development. During development, NPCs switch from initial expanding divisions to neurogenic divisions, which marks the onset of neurogenesis. Here, we aimed to understand when NPCs switch division modes to generate the first neurons in the anterior-most part of the zebrafish brain, the telencephalon. To this end, we used the deep learning-based segmentation method Cellpose and clonal analysis of individual NPCs to assess production of neurons by NPCs in the first 24 hours of zebrafish telencephalon development. Our results provide a quantitative atlas detailing the production of telencephalic neurons and NPC division modes between 14 and 24 hours post-fertilization. We find that within this timeframe, the switch to neurogenesis is gradual, with considerable heterogeneity in individual NPC neurogenic potential and division rates. This quantitative characterization of initial neurogenesis in the zebrafish telencephalon establishes a basis for future studies aimed at illuminating the molecular mechanisms and regulators of early neurogenesis.

## Introduction

During embryonic development of the vertebrate brain, spatially distinct regions with specific cognitive and information-processing functions emerge. Underlying these different functions is the spatiotemporally controlled generation of the different types of neurons present in each brain region (Ge et al., 2022; Lodato and Arlotta, 2015; Shohayeb et al., 2021). At the root of this neuronal diversity are neural stem and progenitor cells (NPCs) that are present within the developing neuroepithelium and undergo cell division to generate new neurons (Ortiz-Álvarez and Spassky, 2021).

In vertebrates, embryonic NPCs typically constitute polarized and elongated cells with apical and basal processes that span the neuroepithelium from the ventricular surface to the meninges. As the cell cycle progresses, the NPC nucleus moves apically in interkinetic nuclear migration. In most vertebrate species, NPCs undergo mitosis exclusively at the surface of the brain ventricle. In initial phases of vertebrate brain development, NPCs undergo symmetric proliferative divisions to expand their pool size. Subsequently, NPCs switch to asymmetric and symmetric neurogenic division modes to mediate the production of the first neurons at the onset of neurogenesis (Casas Gimeno and Paridaen, 2022; Delaunay et al., 2017). Newborn neurons arising from these divisions detach from the ventricular surface and migrate basally to their final position. Over time, different types of neurons are generated from the same initial pool of NPCs as development proceeds.

Despite the advances made in understanding how the production of neuronal diversity from embryonic NPCs is regulated (Ge et al., 2022; Lodato and Arlotta, 2015), our understanding of the molecular and cell biological mechanisms underlying individual NPC division mode and fate decisions is limited. For instance, studies using various NPC lineage-tracing methods have demonstrated that there is remarkable heterogeneity in the division modes used by individual NPCs in the vertebrate nervous system (Dong et al., 2012; Gao et al., 2014; Gomes et al., 2011; He et al., 2012; Hevia et al., 2021; Llorca et al., 2019). Also, it is not clear whether the embryonic NPCs present at the onset of neurogenesis are equipotent in their ability to produce the different types of neurons (Ge et al., 2022; Zechner et al., 2020). In order to understand the factors and processes underlying individual NPC biology in each part of the nervous system, it is important to have a clear picture of the timing of neurogenesis and NPC division mode changes in each brain region.

The most anterior part of the developing neural tube gives rise to the telencephalon, the diencephalon and the retina. In the developing mouse dorsal telencephalon the spatiotemporal pattern of neuron production and the timing of NPC division mode changes are well established. However, the early neurogenic process of the dorsal telencephalon in the zebrafish model has not been described in as much detail. In zebrafish, the dorsal telencephalon constitutes the embryonic primordium of the pallium, which is evolutionarily related to the mammalian neocortex (Diotel et al., 2020; Wullimann, 2009). Seminal studies showed that the first neurons in the zebrafish forebrain are born between 14 and 16 hours post-fertilization (hpf) in two major neuronal clusters, the dorso-rostral and ventro-rostral clusters (Korzh et al., 1993; Macdonald et al., 1994; Ross et al., 1992; Wilson et al., 1990). These clusters correspond to neurons of the prospective telencephalon and ventral diencephalon, respectively. A more recent study using time-lapse imaging of NPCs in the zebrafish forebrain between 26 and 48 hpf provided the first description of NPC division modes after the onset of neurogenesis (Dong et al., 2012). However, the complete time course of individual NPC division mode switches and how this connects to the production of the first neurons in zebrafish pallium development has not been described in detail.

Here, we set out to characterize neuron production and individual NPC lineages during the first day of zebrafish pallium development. Using machine learning-based NPC cell segmentation and counting as well as sparse lineage tracing techniques, we quantified neuron production within the telencephalon. We observed variations in the total cell and neuron counts between individual embryos of the same stage. Inspection of individual NPC behavior showed heterogeneity in division rate and neurogenic potential during initial stages of neurogenesis in the zebrafish telencephalon. This characterization of NPC biology during early neurogenesis of the zebrafish pallium will be instrumental for future in-depth study of the rules governing individual NPCs behavior in establishing the neuronal diversity and complexity of the adult brain.

## Results

### Neurogenesis in the zebrafish telencephalon starts after 14 hpf

To assess the overall morphological changes in the first 24 hours of telencephalon development, we first explored the general tissue architecture of the anterior-most part of the neural tube between 12 and 22 hpf. We examined forebrain morphogenesis by imaging fixed DAPI-stained samples (Fig. 1). As described previously (Affaticati et al., 2015; Werner et al., 2021), the anterior neural tube transitioned from an initially disorganized and unpolarized epithelium into a pseudostratified polarized neural tube-like structure (Fig. 1A) with formation of optic vesicles between 14 and 18 hpf. As described previously (Korzh et al., 1993; Macdonald et al., 1994; Ross et al., 1992; Wilson et al., 1990), by 22 hpf neurons were present within the telencephalon (as marked by the neuronal-specific small RNA-binding proteins HuC/D) (Fig. 1B).

**Figure 1.**
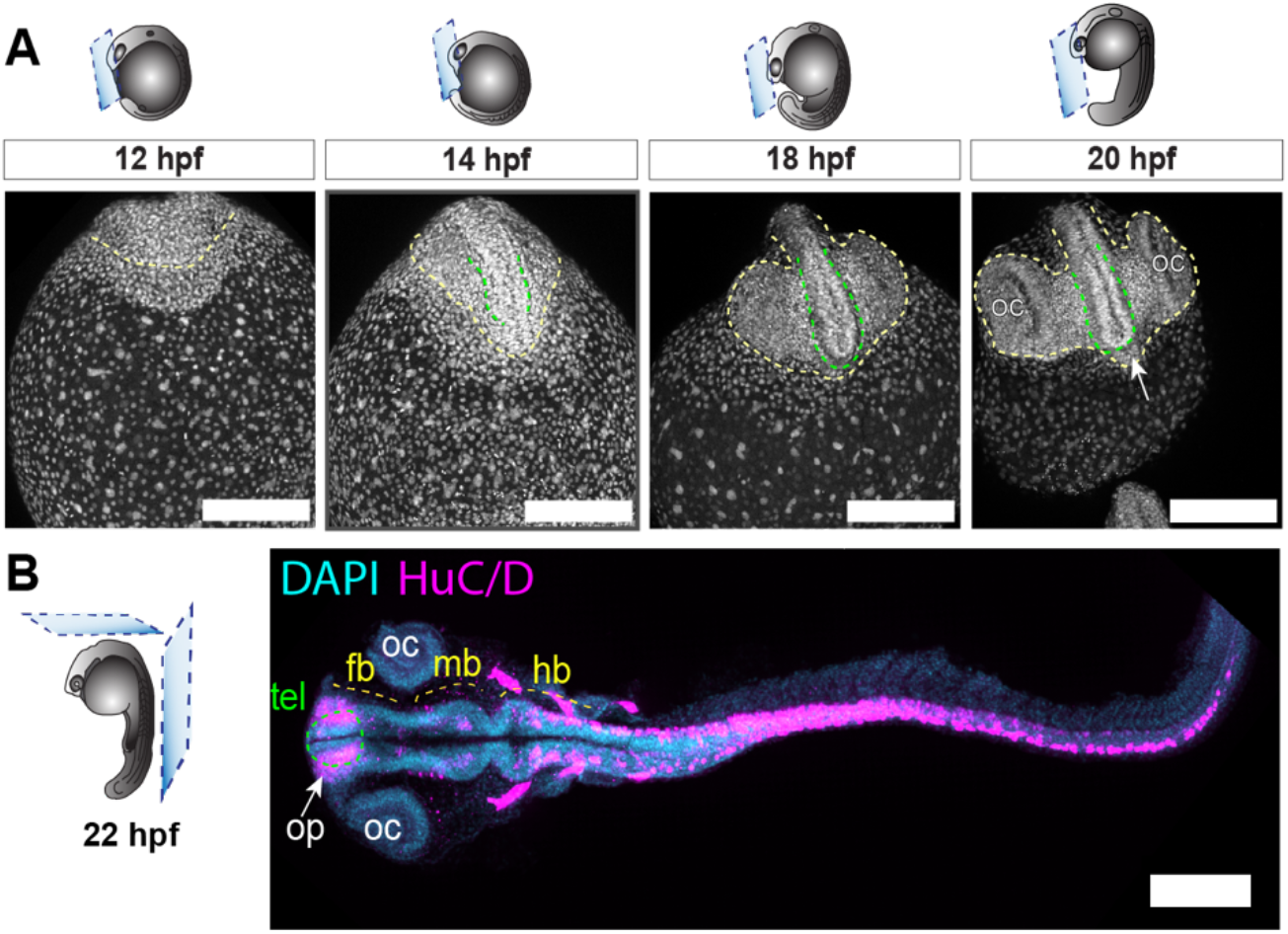
Telencephalon development in zebrafish embryogenesis. A) Frontal views of the anterior-most part of zebrafish embryo heads stained with DAPI to label nuclei at the indicated timepoints. Dorsal is up. The developing neural territory is delineated by a yellow dashed line and the developing telencephalon is highlighted by a green dashed line. At 20 hpf, the structures labelled are OC: optic cup, OP: olfactory placode, arrow: hypothalamus. Scale bar is 200 μm. B) Dorsal view of a zebrafish embryo (flat-mounted) at 22 hpf stained with DAPI to label nuclei and the pan-neuronal marker HuC/D (magenta). Anterior is to the left. The major segments of the CNS are highlighted with a yellow dash line. The telencephalon (Tel) is delineated with a green dashed line. Scale bar is 150 μm. FB, forebrain; hpf: hours post-fertilization; MB, midbrain; HB, hindbrain; SC, spinal cord; OC, optic cup; OP, olfactory placode.

Next, we characterized the emergence of neurons during the first 24 hours of telencephalon development in more detail. At 14 hpf, the neuroepithelial cells in the telencephalon did not show clear apicobasal polarity and no defined ventricular lumen was visible. In contrast, at 16 hpf the nuclei showed radial alignment indicative of pseudostratified epithelial architecture (Fig. 2A, first and second column) and the ventricle was now visible at the midline. Immunofluorescent staining for HuC/D showed that the first neurons within the telencephalon were born between 14 hpf and 16 hpf (Figure 2A, first and third row). At 16 hpf, the first neurons were observed in the middle section of the dorsoventral axis (Fig. 2A). This confirms the timing of initial neurogenesis as originally reported (Ross et al., 1992). As in other developing neuroepithelia, the neuronal cell bodies are localized basally, away from the ventricular surface, while progenitor cells form a defined germinal zone on the apical side of the neural epithelium. At 16 hpf, many newborn neurons maintained apico-basal connections (Fig. 2B), indicating that newborn neurons maintain the apicobasal morphology for some time after mitosis and before detachment of the apical process (Alexandre et al., 2010).

**Figure 2.**
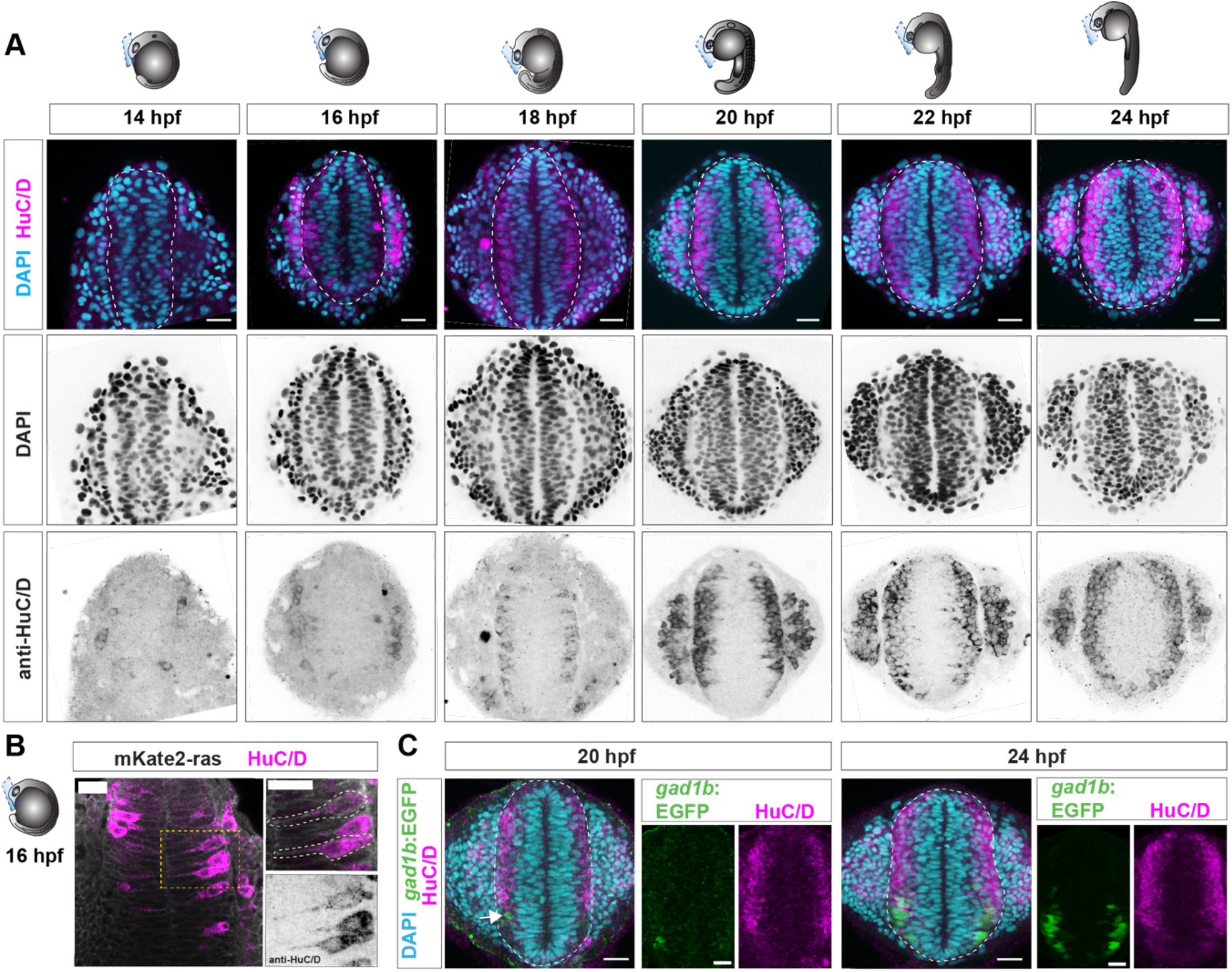
Neurogenesis in the developing telencephalon. A) Representative (single-optical section) confocal images of the frontal view of the telencephalon (outlined with dashed line) of zebrafish embryos between 14 and 24 hpf stained with DAPI to label nuclei (cyan) and immunostained with anti-HuC/D to label neurons (magenta). Dorsal is up. Top row shows the merged DAPI (cyan), HuC/D (magenta) channels; middle row shows DAPI nuclei signal in inverted gray and bottom row shows HuC/D fluorescence signal in inverted gray. Scale bar is 25 μm. B) Frontal view (single-optical confocal section) of the telencephalon of a 16 hpf embryo immunostained against the pan-neuronal marker HuC/D (magenta) and expressing membrane-targeted mKate2 (grey). The magnification images show HuC/D+ neurons that have an apical attachment. Scale bar is 20 um. C) Representative (single-optical section) confocal images of frontal view of the telencephalon (dashed line) showing tg(*gad1b*:EGFP)+ GABA-ergic neurons (green), nuclei (DAPI, cyan) and pan-neuron label (HuC/D, magenta) at 20 hpf (left) and 24 hpf (right). The arrow indicates an individual GABA-ergic neuron. Scalebar is 25 μm. hpf, hours post-fertilization.

The adult telencephalon is subdivided into the dorsal-most pallium, containing mainly glutamatergic excitatory neurons and the ventral subpallium, that harbors mainly GABAergic interneurons (Than-Trong and Bally-Cuif, 2015; Wullimann, 2009). Previous studies described GABAergic neurons in the telencephalon from 27 hpf onwards (Macdonald et al., 1994; Martin et al., 1998). In order to assess the timing of the onset of GABAergic neurogenesis in the telencephalon more precisely, we assessed the presence of GABAergic neurons using the *gad1b*::EGFP transgenic line (Satou et al., 2012). Examination of EGFP expression between 16 and 24 hpf showed that a low number of GABAergic neurons were present from 20 hpf onwards (Fig. 2C). At 24 hpf, the ventral-most section of the telencephalon neuronal layer contained GABAergic neurons. This shows that soon after the onset of telencephalic neurogenesis, distinct neuronal cell fates are specified that emerge in spatially distinct domains. This positional neuronal diversity probably depends on dorsoventral morphogen gradients similar to that in the spinal cord (Ge et al., 2022; Sagner and Briscoe, 2019).

### Neuron production increases after 18 hpf in the telencephalon

In order to quantify the total numbers of progenitors and neurons at the first stages of telencephalon development, we made use of Cellpose, a recently published machine-learning-based cellular segmentation algorithm (Stringer et al., 2021). We set up an analysis pipeline to count nuclei and neurons within the telencephalon (Fig. 3A; see Methods for details). After preprocessing the raw image data to correct for changes in signal brightness due to imaging depth, we specified the boundaries of the telencephalon for each set of 3D microscope images (Fig. 3A, Suppl. Figs 1, 2). Using the nuclear staining signal, the Cellpose algorithm was used to segment individual cell nuclei within the 3D image stack. To obtain total cell numbers within the telencephalon (Fig. 3B), the number of unique nuclear labels was counted. We then used the HuC/D immunofluorescent signal to assign neuronal identities within the telencephalon region for each set of 3D images (Fig. 3A). To obtain total neuron numbers, the number of cells assigned neuronal identity within the telencephalon was counted (Fig. 3A, D). Assuming the absence of cell death (Cole and Ross, 2001) and absence of other cell types such as microglia (Herbomel et al., 2001) in the telencephalon prior to 24 hpf, we assigned NPC identity to all non-HuC/D positive cells. In this way, we obtained total cell number counts, total neuronal counts and total NPC counts within the telencephalon at the embryonic stages of 14 to 24 hpf (Fig. 3B-F, Table 1, Suppl. Fig. 3, Suppl. Table 1).

**Figure 3.**
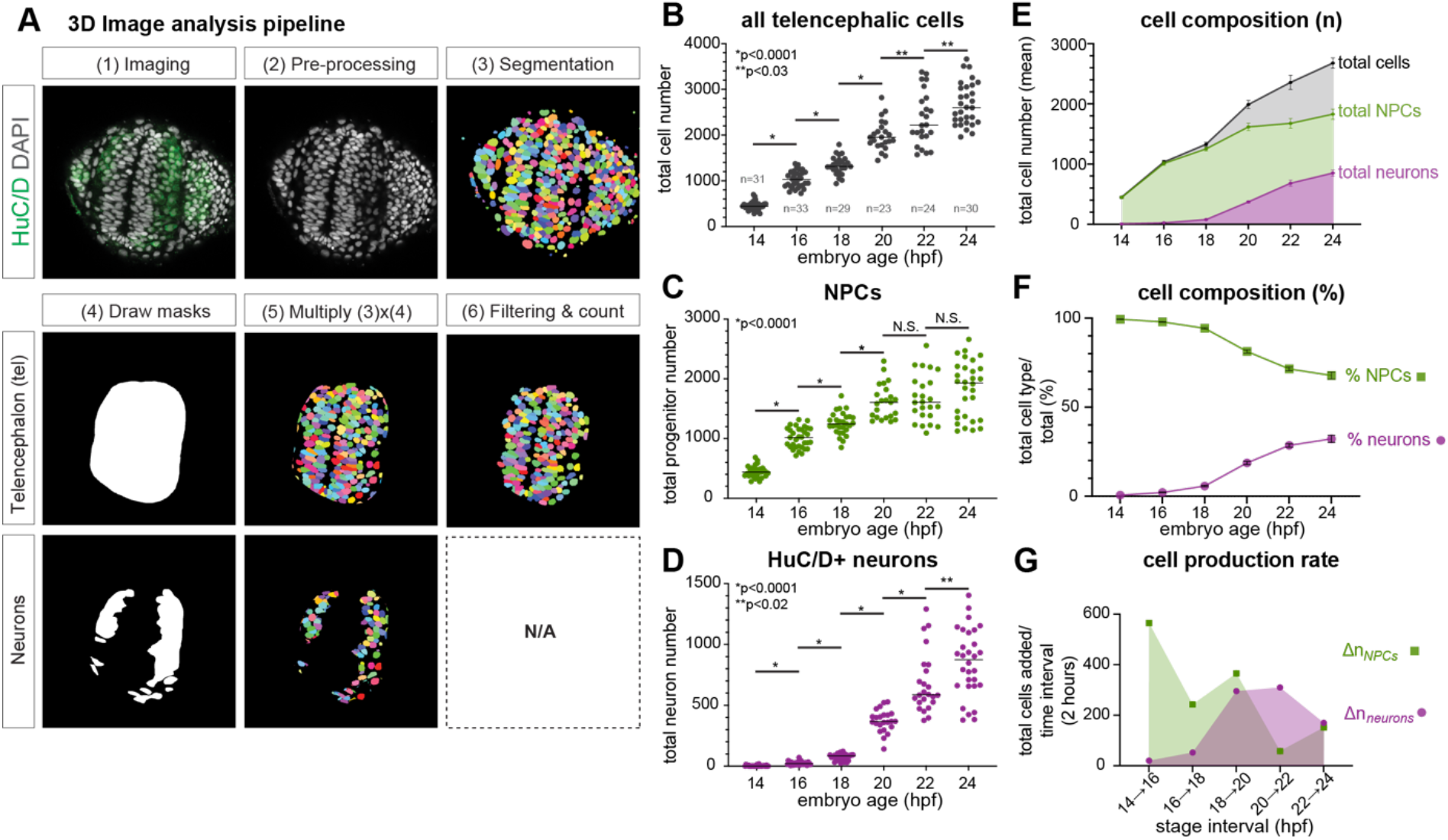
Quantification of cell numbers in the developing telencephalon. A) Pipeline of the steps towards quantification of total cell number and neuronal number using Cellpose. Further detail can be found in material and methods. B-D) Quantification of the total number of cells (B), NPCs (C) and D) neurons present in the telencephalon between 14 and 24 hpf as determined using the Cellpose pipeline (data from N=3-4 experiments with n of embryos/stage indicated). Each point reflects data from one individual embryo. P-values are indicated for comparison between subsequent stages (two-tailed Mann-Whitney test). E-F) Cell composition (E) and percentage (F) of NPCs (green) and neurons (magenta) between 14 and 24 hpf. Data are mean ± SEM. G) Cell production rate (total cells added per 2-hour time- interval) for NPCs (green) and neurons (magenta) between 14 and 24 hpf. hpf, hours post-fertilization; tel, telencephalon.

**Table 1.**
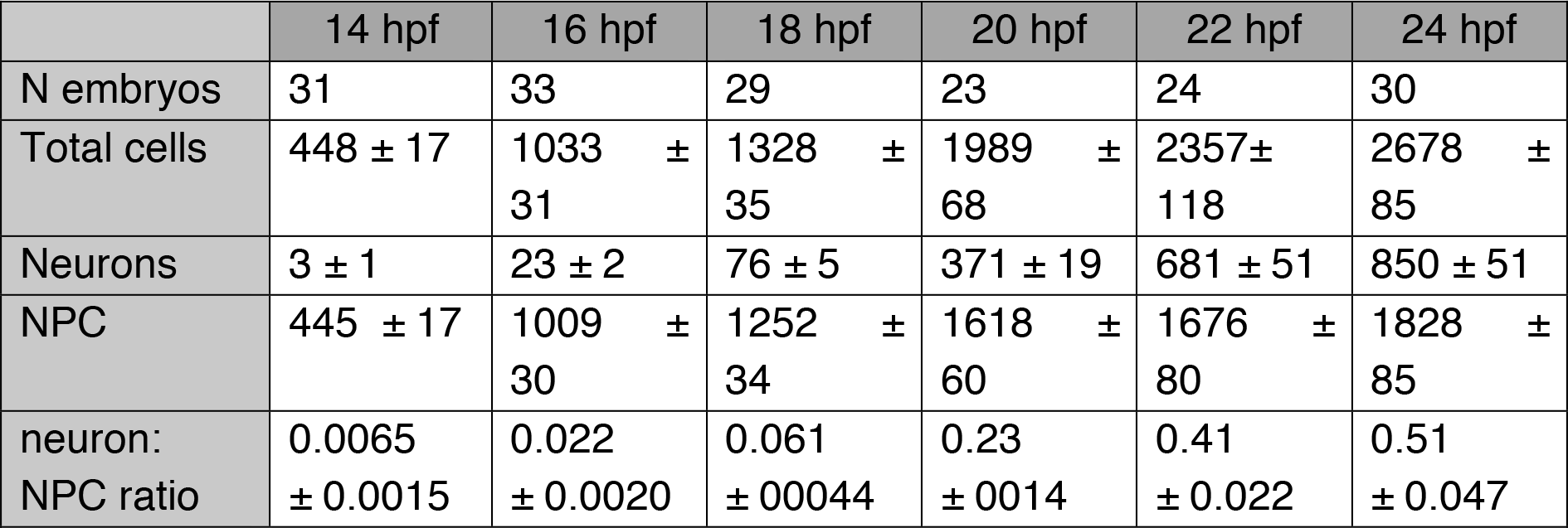
Summary of total cell, neuron and NPC number in the telencephalon between 14 and 24 hpf. Average and standard error of the mean of the number of total cells, neurons and NPCs (calculated as n_total cells_ – n_neurons_) counted from the indicated number of embryos at the indicated timepoints. The neuron:NPC ratio was calculated for each individual embryo.

|  | 14 hpf | 16 hpf | 18 hpf | 20 hpf | 22 hpf | 24 hpf |
| --- | --- | --- | --- | --- | --- | --- |
| N embryos | 31 | 33 | 29 | 23 | 24 | 30 |
| Total cells | 448 ± 17 | 1033 ± 31 | 1328 ± 35 | 1989 ± 68 | 2357 ± 118 | 2678 ± 85 |
| Neurons | 3 ± 1 | 23 ± 2 | 76 ± 5 | 371 ± 19 | 681 ± 51 | 850 ± 51 |
| NPC | 445 ± 17 | 1009 ± 30 | 1252 ± 34 | 1618 ± 60 | 1676 ± 80 | 1828 ± 85 |
| neuron:<br>NPC ratio | 0.0065<br>± 0.0015 | 0.022<br>± 0.0020 | 0.061<br>± 0.0044 | 0.23<br>± 0.014 | 0.41<br>± 0.022 | 0.51<br>± 0.047 |

Overall, the absolute number of total cells showed a linear increase between 14 and 24 hpf (Fig. 3B). The number of NPCs was substantially increasing between 14 and 20 hpf, after which it reached a plateau (Fig. 3C). In contrast, the first neurons emerged between 14 and 16 hpf (Fig. 3D). After 18 hpf, the number of neurons was rapidly increasing. By 24 hpf, about one third of the total cells in the telencephalon are neurons (Fig. 3E, F). Strikingly, there was considerable variation in the total cell and total neuron number counted in the telencephalon of individual embryos (Suppl. Fig. 3; see also discussion), especially at the 22 and 24 hpf stages. The coefficient of variation (CV) was particularly larger for the neuron counts than the total cell counts (Suppl. Fig. 3A, B; 22 hpf: CV 36.4% for neurons *versus* CV 24.6% total cell count; 24 hpf: CV 32.6% neurons *versus* CV 17.5% total cell count). Moreover, there was no strong connection between the total cell and neuron number per embryo (Suppl. Fig. 3C), suggesting that the variations do not stem from embryos simply containing more cells.

Together, these data show that the first neurons are born between 14 and 16 hpf and that neurogenesis in the telencephalon rapidly increases after 18 hpf.

### The division and neurogenic rates change during the first day of telencephalon development

Next, we analysed the absolute number of progenitors and neurons added per time window in order to distill information regarding NPC division and neuron production rates during the first day of telencephalon development (Table 2-3).

**Table 2.** Estimated number of added cells in the telencephalon for 2-hour intervals between 14 and 24 hpf. The number of added cells, NPCs and neurons for each interval (t-1) to (t) was calculated by (n_totalcells(t)_ – n_totalcells(t-1)_), (n_NPC(t)_ – n_NPC(t-1)_) and (n_neurons(t)_ – n_neurons(t-1)_), respectively. The growth rate for each interval (t-1) to (t) was calculated as (n_totalcells(t)_: n_totalcells(t-1)_). The division rate for each interval (t-1) to (t) was calculated by (n_total cells added(t)_ / n_NPC(t-1)_) and neurogenic fraction by (n_neurons added(t)_ / n_total cells added(t)_). Note * corresponds to division rate values that exceed one division per NPC between (t-1) and (t).

|  | 14 to 16 hpf | 16 to 18 hpf | 18 to 20 hpf | 20 to 22 hpf | 22 to 24 hpf |
| --- | --- | --- | --- | --- | --- |
| Total cells added | 584 | 296 | 661 | 368 | 321 |
| NPCs added | 564 | 243 | 366 | 58 | 151 |
| Neurons added | 20 | 53 | 295 | 310 | 169 |
| Growth rate | 2.3 | 1.3 | 1.5 | 1.2 | 1.1 |
| Division rate | 1,3* | 0,29 | 0,53 | 0,23 | 0,19 |
| Neurogenic fraction | 0,034 | 0,18 | 0,45 | 0,84 | 0,53 |

**Table 3.** Estimated number of added cells in the telencephalon between 4-hour intervals between 16 and 24 hpf. The number of added cells, NPCs and neurons for each interval (t-1) to (t) was calculated by (n_totalcells(t)_ – n_totalcells(t-1)_), (n_NPC(t)_ – n_NPC(t-1)_) and (n_neurons(t)_ – n_neurons(t-1)_), respectively. The division rate for each interval (t-1) to (t) was calculated by (n_total cells added(t)_ / n_NPC(t-1)_) and neurogenic fraction by (n_neurons added(t)_ / n_total cells added(t)_).

|  | 16 to 20 hpf | 20 to 24 hpf |
| --- | --- | --- |
| Total cells added | 957 | 689 |
| NPCs added | 609 | 209 |
| Neurons added | 348 | 479 |
| Division rate | 0,95 | 0,43 |
| Neurogenic fraction | 0,36 | 0,70 |

Between 14 and 16 hpf, there is a large increase in cell number and the division rate exceeds one division per NPC (Fig. 3B, Table 2). Based on the cell cycle length reported for zebrafish segmentation stages of ± 4 hours (Kimmel et al., 1994), we assume that during the two-hour interval a single NPC would not divide more than once. Therefore, the increase in total cells probably reflects the addition of new NPCs through the extensive neurulation cell movements that are still taking place prior to 16 hpf (Ivanovitch et al., 2013; Werner et al., 2021).

After 16 hpf, neurulation movements have seized and new cells added are therefore likely to be generated from NPC divisions. After 16 hpf, the total cell number showed an almost linear increase (Fig. 3E-G; Table 2). However, after 20 hpf, the NPC division rate was lower compared to before 20 hpf (Table 2-3). This could be due to increased NPC cell cycle length or a reduction in the number of NPCs that are actively proliferating after 20 hpf. In contrast, the fraction of newly generated cells that were neurons (neurogenic fraction) increased after 20 hpf (Table 2-3), indicating a relative increase in the production of neurons per NPC.

We quantified the number of cells, NPCs and neurons within the telencephalon. Thus, the ratio between number of newborn NPCs and neurons provides insight into the balance between the generation of NPC and neuronal daughter cells from the pool of NPCs. For instance, for the 18→20 hpf interval, added NPCs and neurons constitute approximately equal proportions. Comparison of the total of added NPCs *versus* neurons at each 2-hour interval between 16 and 24 hpf (Fig. 3G) showed that prior to 18 hpf, there are proportionally more NPCs added than neurons, while after 22 hpf more neurons are added. This switch is likely to reflect overall switches in NPC division mode from proliferative (generating new NPCs) to neurogenic (generating one or two neurons).

### Clonal analysis of individual neural progenitor lineages in the first day of telencephalon development

In order to obtain information on division modes and neuron production by individual NPCs, we next performed clonal analysis of individual telencephalic NPCs using the Zebrabow system (Pan et al., 2013). New NPCs are born from symmetric proliferative divisions. Neurons are born from two types of neurogenic divisions. In asymmetric (P-N) division, a new NPC and a neuron are generated. In symmetric terminal (N-N) divisions, the mother NPC is consumed while two neurons are generated. To induce sparse labelling of individual progenitors allowing exploration of their progeny, we induced recombination by injection of *Cre^ERT2^* mRNA and activation of Cre-mediated Zebrabow cassette recombination by 4-OHT treatment from 10 to 11 hpf, just prior to the onset of neurogenesis (Fig. 4A). First, we titrated the amount of injected *Cre^ERT2^* mRNA to limit instances of recombination in the absence of 4-OHT treatment (Suppl. Fig 4A). Next, we generated sparse Zebrabow clonal labelling by mosaic injections of *Cre^ERT2^* mRNA into Zebrabow embryos, followed by 4-OHT treatment at 10 hpf, and analysis of the resulting clones in the telencephalon at 16, 20 or 24 hpf. To facilitate identification of clone size and composition, mRNA encoding membrane-targeted fluorescent protein (LSSmOrange-ras) was coinjected and HuC/D immunofluorescence used to readout neuronal fate (Fig. 4B). For each individual clone (LSSmOrange+ and with the same Zebrabow colour combination), the clone size (cells generated from a single NPC) and composition was determined (Fig. 4B, C; Suppl. Fig. 5-D; Suppl. Table 2).

**Figure 4.**
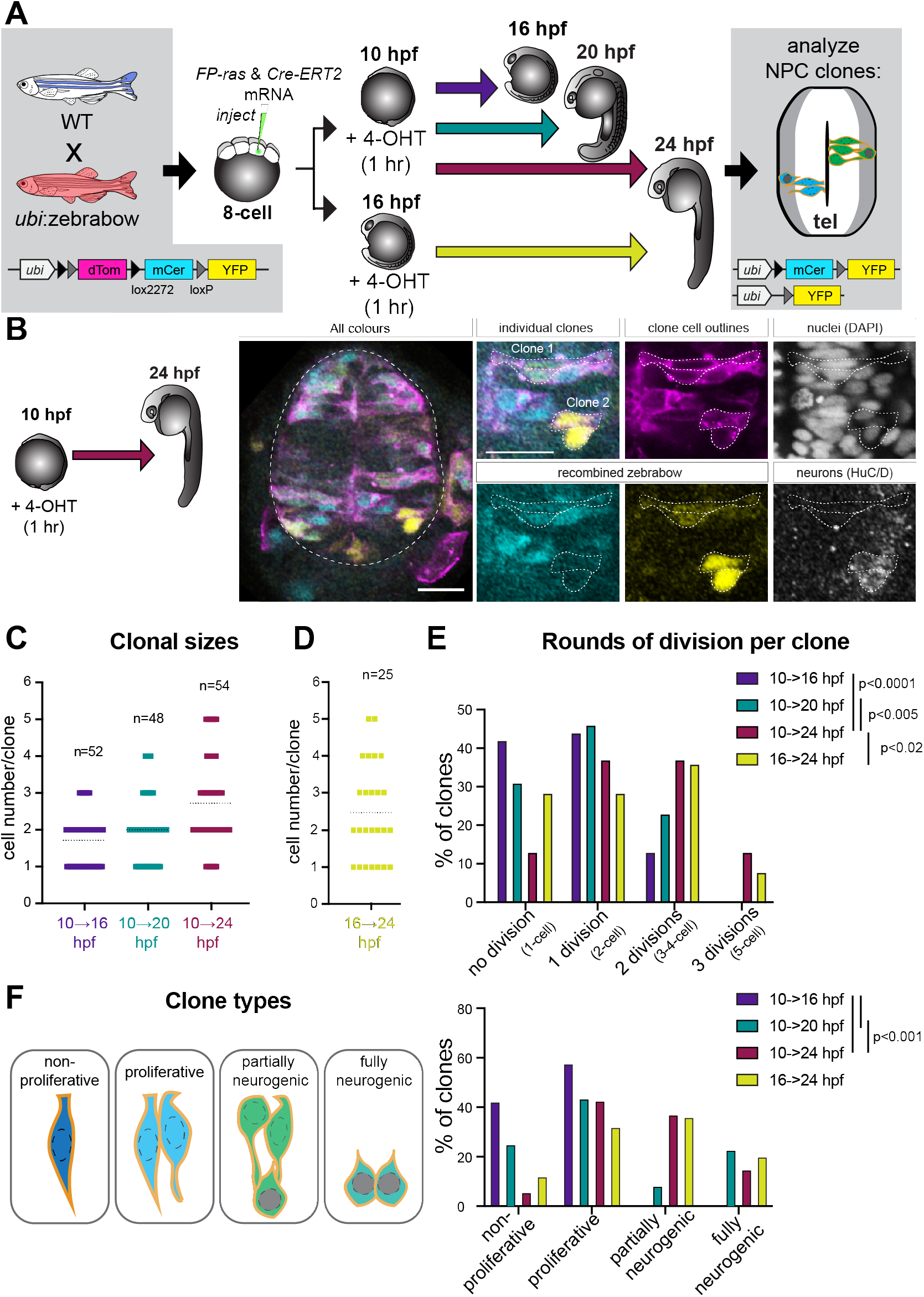
Clonal analysis of NPCs in the developing telencephalon. A) Experimental design of clonal labeling of NPCs by injection of CreERT2 mRNA into *ubi*:zebrabow embryos, followed by induction of recombination through 4-OHT treatment and analysis at the indicated stages of the resulting NPC clones. B) Representative (single-optical section) confocal image of the frontal view (membrane-targeted LSSmOrange-ras (magenta); recombined zebrabow cassette (mCer, cyan; EYFP; yellow)) of the telencephalon (outlined with dashed line) of *ubi*:zebrabow embryos recombined at 10 hpf and investigated at 24 hpf. Dorsal is up. The magnifications show individual fluorescent channels for membrane-targeted LSSmOrange-ras (magenta), nuclei (DAPI; grey), recombined zebrabow cassette (mCer, cyan; EYFP; yellow) and the pan-neuronal marker HuC/D (grey) for two separate NPC clones. The outline of each clone is indicated by dashed lines. Scale bar is 25 μm. C, D) Clone size distributions for clones induced at 10 hpf (C) or 16 hpf (D) and analyzed at 16, 20 or 24 hpf. The mean clone size is indicated by the dashed line. N of clones is indicated for each induction + chase group. E) The frequency of clones with distinct number of cell division rounds taking place in that clones’ lineage for each induction + chase group. P-values are indicated for the comparison between induction + chase groups (Fisher’s exact test). F) The frequency of different clone types (non-proliferative, proliferative (only NPC), partially neurogenic (NPC+neuron) and fully neurogenic (only neuron)) for each induction + chase group. P-values are indicated (Fisher’s exact test). hpf, hours post-fertilization.

Comparison of clones induced at 10 hpf and analysed at 16, 20 and 24 hpf, respectively, showed an increase in average clone size and the range of clone sizes over time (Fig. 4C). Similarly to what was observed for individual NPC clones in the developing zebrafish retina (He et al., 2012), there was considerable heterogeneity in clone size (Fig. 4C) and the estimated number of cell divisions (Fig. 4E; Suppl. Fig. 5A) for each clone. Overall, the proportion of one-cell-sized clones was decreasing when comparing clones at 16, 20 and 24 hpf stages, showing that the vast majority of individual NPCs underwent at least one cell division between 10 and 24 hpf.

In line with the low number of neurons observed within the entire telencephalon at 16 hpf, no clones were observed containing neurons at that stage (Fig. 4F). At 20 hpf, most individual clones were either proliferative (lacking neurons) or fully neurogenic (only neurons). This suggests that only a subset of individual NPCs are undergoing symmetric neurogenic divisions at this time and that these constitute mainly terminal divisions in which the mother NPC is lost. In contrast, at 24 hpf, we observed proliferative, partially neurogenic and fully neurogenic clones (Fig. 4F).

Overall, these observations indicate that the onset of neurogenesis and the division rate is not equal for all individual NPCs and that they gradually switch to neurogenesis between 16 and 24 hpf.

### The growth rate and neuron production per progenitor are comparable between total telencephalon and clonal analysis approaches

Next, we aimed to directly compare our Cellpose-based quantifications and clonal analyses. To avoid the influence of new NPCs being added to the telencephalon region by neurulation movements, we compared Cellpose-based quantifications between 16 and 24 hpf with those based on zebrabow clones induced at 16 hpf and analyzed at 24 hpf (Fig. 4D-F, Table 4, Suppl. Fig. 4B, C; Suppl. Fig. 5E; Suppl. Table 2). We observed that the average growth rate based on average clone size and total cell and NPC counts was in the same range (2.6x telencephalon growth rate from 16 to 24 hpf with Cellpose *versus* 2.4x average clone size 16 to 24 hpf Zebrabow). When comparing the average number of neurons generated by 24 hpf from one single NPC at 16 hpf, these numbers were again similar (0.84 neuron/NPC Cellpose; 0.72 neuron/NPC Zebrabow). When combining all clones together, the proportion of neurons at 24 hpf was also very similar to the Cellpose-based quantification (32.2% Cellpose; 30.6% 10→24 hpf Zebrabow; 29.0% 16→24 hpf Zebrabow).

**Table 4.** Comparison of cell and neuron production per NPC from Cellpose- and Zebrabow-based analyses.

|  | 10 to 16 hpf<br>Zebrabow | 10 to 20 hpf<br>Zebrabow | 10 to 24 hpf<br>Zebrabow | 16 to 24 hpf<br>Zebrabow | 16 to 24 hpf<br>Cellpose |
| --- | --- | --- | --- | --- | --- |
| Average clone size | $1.7 \pm 0.10$ | $2.0 \pm 0.13$ | $2.7 \pm 0.17$ | $2.4 \pm 0.26$ | |
| Number of cells per initial NPC<br>($n_{\text{totalcells}(24\text{hpf})} / n_{\text{NPC}(16\text{hpf})}$) | | | | | 2.6 |
| Average number neurons/clone | 0 | $0.48 \pm 0.11$ | $0.83 \pm 0.14$ | $0.72 \pm 0.15$ | |
| Number of neurons per initial NPC<br>($n_{\text{neurons}(24\text{hpf})} / n_{\text{NPC}(16\text{hpf})}$) | | | | | 0.84 |

Therefore, we conclude that the average NPC clonal size and composition is representative of neuron production within the entire telencephalic brain region.

### Neural progenitor cells change division mode around 20 hpf

Next, we explored whether the data we obtained can inform on the division modes used by individual NPCs between 16 and 24 hpf (Fig. 5A). As the Cellpose-based quantifications estimates the absolute number of NPCs present within the telencephalon, we assume that once neurulation movements have finished prior to 16 hpf as previously described (Affaticati et al., 2015; Werner et al., 2021), the cells born within a certain time frame are progeny of the pool of NPCs present at the onset. Using this assumption, we calculated the estimated numbers of total division events, symmetric proliferative, asymmetric and symmetric neurogenic divisions within the telencephalon for several stage intervals (Fig. 5B, C). Over progressive time intervals, the relative proportion of symmetric proliferative (P-P divisions) division were decreasing, while the proportion of asymmetric (P-N division) and symmetric terminal (N-N divisions) division events were increasing after 18 hpf. Between 16 and 18 hpf, P-P division events were at least 8-fold more likely than P-N and N-N divisions. After 18 hpf, the separation between the range of P-P, P-N and N-N division events was progressively disappearing, suggesting that the probability of P-P was nearing that of P-N and N-N divisions (Fig. 5C).

**Figure 5.**
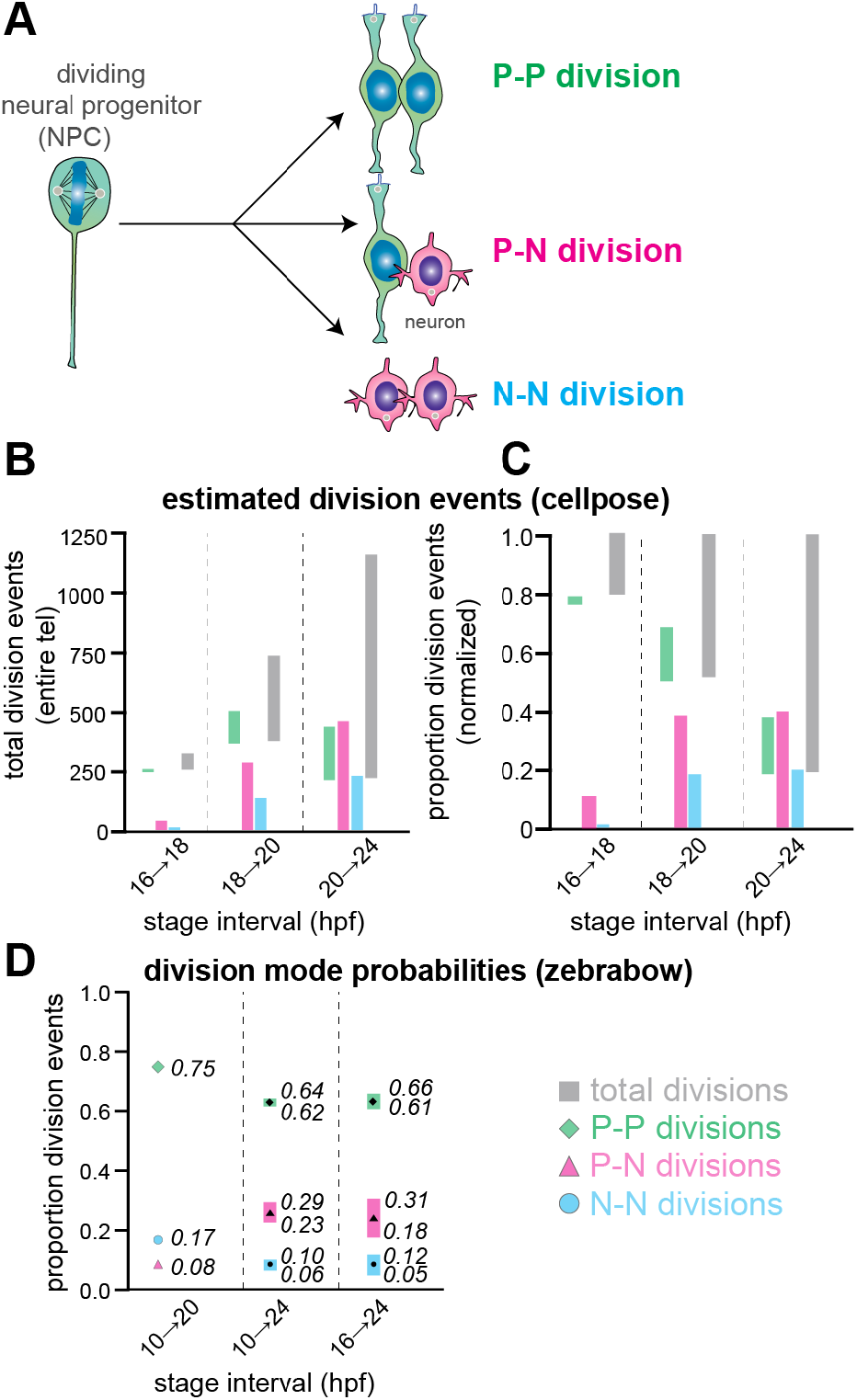
Estimated number of division events and division mode probabilities during onset of telencephalic neurogenesis. A) Dividing NPCs have three possible division modes: symmetric proliferative (P-P; green), asymmetric neurogenic (P-N; magenta) and symmetric neurogenic (N-N; cyan) divisions. B, C) Range of estimated absolute (B) or normalized (C; to maximum estimate of all division events) division events for each division mode type based on Cellpose-based quantifications for the stage interval indicated. Grey bars indicate the min-max range of estimated total cell division events. D) Division mode probabilities for each division mode type based on Zebrabow-based quantifications of the indicated induction + chase groups. For induction at 10 and chase to 20 hpf, the probabilities are a single value, and for 10 to 24, and 16 to 24 hpf, the min-max probabilities are shown. hpf, hours post-fertilization.

Next, we calculated the probabilities for each division type based on the zebrabow clonal analysis. Here, for each clone, we determined which division modes were theoretically used within its lineage leading up to that clones’ particular size and composition. We then used this information to estimate the probability for each division mode per division event in the history of that clone (Fig. 5D). Prior to 20 hpf, the probability for asymmetric division was low (0.08), while for the 10 to 24 and 16 to 24 hpf clonal analysis, it had increased to ≈0.25 (Fig. 5D). Strikingly, N-N division probability was higher prior to 20 hpf compared to between 20 and 24 hpf. The probability of P-P divisions was higher for chase period up to 20 hpf compared to chase periods up to 24 hpf, indicating that after 20 hpf, P-P division probability is decreasing as suggested also by the Cellpose-based division event estimates.

Taken together, these results show that the onset of neurogenesis is accompanied with an initial switch around 16 hpf of a subset of NPCs from symmetric proliferative to symmetric neurogenic divisions, followed by another switch around 20 hpf to an asymmetric division mode.

## Discussion

To understand how stem and progenitor cells generate the complex structures of adult organisms, it is key to understand progenitor biology and regulation thereof at the individual cell level. *In vivo* studies in invertebrate and vertebrate models have elucidated some of the molecular and cellular mechanisms, such as sequential expression of transcription factors (Pollington et al., 2022), acting in progenitor division mode selection. However, to study individual progenitors in a specific tissue or regions, it is key to have an in-depth understanding of the developmental timeline of differentiation within that tissue.

The onset of neurogenesis and overall NPC division mode changes have been well described for areas of the zebrafish nervous system such as the hindbrain, spinal cord and retina (Boije et al., 2015; He et al., 2012; Hevia et al., 2021; Kimura et al., 2008; Nerli et al., 2022; Satou et al., 2012). In this experimental work, we extend the existing body of work with an quantitative characterization of the first wave of neurogenesis in the developing zebrafish telencephalon using machine-learning based nuclear segmentation and clonal analysis methods. Our results on the timing of neurogenesis match that of seminal earlier studies (Korzh et al., 1993; Ross et al., 1992) and confirm that the first primary neurons in the zebrafish telencephalon are born between 14 and 16 hpf. Moreover, we extend earlier studies by quantifying the cells present in the telencephalon prior to 24 hpf using the machine-learning based nuclear segmentation algorithm Cellpose (Stringer et al., 2021). This allows to extrapolate information about neuron production rates and NPC division modes that is valuable for follow-up studies aimed at understanding NPC biology at the single-cell level.

We observed considerable variability in total cell, NPC and neuron counts that we obtained for individual embryos of the same stages, especially after 20 hpf. Part of this variation may be due to differences between embryo clutches and technical limitations such as inconsistencies in setting the 3D boundaries of the telencephalon and weakening of the fluorescent signal deeper into the tissue as the telencephalon is growing after 20 hpf. However, the variation in total neuron numbers between individual embryos of the same stage may have a biological component. Strikingly, a recent study looking at a sub-population of functionally distinct endoderm cells in early zebrafish embryos found similar fluctuations in the total cell number between individual embryos (Moreno-Ayala et al., 2021). This suggests that individual fluctuations in the absolute number of specialized cells within a tissue is a common phenomenon in early (zebrafish) development.

Our key result is the quantification of neurogenesis at the tissue level along the first hours of telencephalon development. We find that NPCs in the telencephalon first expand through proliferative divisions and later there is a balance between neurogenic and proliferative divisions. This is consistent with the general sequence of division modes found in other models of neurogenesis such as the mouse developing neocortex (Gao et al., 2014; Llorca et al., 2019). Our characterization of clonal behavior of individual NPCs showed heterogeneity of clonal trajectories similar to other regions of the zebrafish nervous system (He et al., 2012; Hevia et al., 2021). A previously published clonal analysis of zebrafish forebrain NPCs between 26 and 37 hpf showed division mode probabilities in line with our observations (0.55 P-P divisions; 0.38 P-N divisions; 0.08 N-N division (Dong et al., 2012)). Thus, combining these observations, this suggests that between 16 and 24 hpf, NPCs are gradually switching to neurogenic divisions and that the probability of P-P divisions gradually decreases, while the P-N division mode probability is slowly increasing towards 37 hpf. Strikingly, after an initial peak between 14 and 20 hpf in N-N divisions, the probability for N-N divisions remains low (<0.1) until 37 hpf (our observations and (Dong et al., 2012). Our results indicate that within the telencephalon, proliferative and neurogenic divisions coexist in time, opening the possibility that sub-populations of NPCs show differential neurogenic competence. Furthermore, the presence of different neuronal type domains by 24 hpf indicates that similarly to the spinal cord (Sagner and Briscoe, 2019), differential signaling gradients may exist to mediate distinct neuronal fates born from otherwise similar NPCs.

The dorsal telencephalon gives rise to the pallium, the structure that has been identified as the homologous structure to the mammalian neocortex. Previous studies using clonal analysis methods in the embryonic mouse developing neocortex have elucidated the timing of neurogenesis onset and division mode switches. Moreover, such studies have demonstrated that individual neural progenitor undergo probabilistic and deterministic decisions in generating neuron and glial cell diversity (Gao et al., 2014; Llorca et al., 2019). Similar studies have been performed in zebrafish hindbrain and retina to elucidate the rules and molecular regulators governing NPC lineage decisions and how NPC competence to generate diverse cell types changes as development proceeds (Boije et al., 2015; Hevia et al., 2021; Nerli et al., 2022; Zhao et al., 2021). Moreover, studies using live imaging microscopy can provide valuable insights into the mechanisms underlying asymmetric versus symmetric division modes (Clark et al., 2021; Kressmann et al., 2015; Nerli et al., 2020).

Our results form the foundation for future studies aimed at further illuminating whether zebrafish telencephalon NPCs differ in their clonal output and how their individual characteristics such as transcriptome, position within the tissue and fate determinant inheritance influences their production of neurons. Furthermore, the application of deep learning-based methods such as Cellpose for determining absolute cell numbers in developing tissues opens avenues for understanding how individual variation can be reconciled with developmental robustness.

## MATERIALS AND METHODS

### Zebrafish lines

Adult zebrafish were kept on a 14h-10h light-dark cycle. Embryos were incubated at 28.5 °C in E3 medium until collection and were staged according to (Kimmel et al., 1995). The following lines were used in this study: AB wild-type, Casper (White et al., 2008), Tg(*vglut2a*::loxP-dsRed-loxP-GFP;*gad1b*::GFP) (Satou et al., 2012); tg(*ub*i::Zebrabow) (Pan et al., 2013); Tg(*nestin*:EGFP) (Lam et al., 2009). All animal work was performed according to European and Dutch Animal Welfare legislation.

### Plasmids

pCS2-mKate2-ras was a gift from Caren Norden (Addgene #105938) (Weber et al., 2014). pCS2-LSSmOrange-ras was generated by replacing mKate2 with LSSmOrange coding sequence (Addgene #37131). pCS2-Cre-ERT2 was generated by inserting PCR-generated ERT2 sequence (with pBigTB-CreERT2 (Addgene #149434) as template) into pCS2-zfCre (Addgene #61391). All plasmids were verified by Sanger sequencing.

### mRNA *in vitro* synthesis

pCS2 vectors, which contain a SP6 promoter in front of the coding sequences of interest, were linearized with appropriate restriction enzymes and purified. The linearized plasmids were used as template for *in vitro* mRNA synthesis using mMESSAGE mMACHINE™ SP6 kit (Thermofisher) following the manufacturer’s protocols. The transcription reaction consisted of 1μg of template DNA, 2μl of 10x Reaction Buffer, 10μl off 2x NTP/CAP, 2μl of enzyme mix and water up to 20μl. Transcription reactions were incubated in thermocycler at 37°C for 2 hours, after which 1μl of TURBO DNase was added and incubated for a further 15 minutes to remove the DNA template. mRNA recovery was done by lithium chloride precipitation. mRNA was resuspended in RNAse-free H2O and adjusted to a concentration of 500 ng/μl. The mRNA solution was then distribute in 1ul aliquots and stored at –80°C until use.

### Zebrafish embryo microinjection of mRNA

For ubiquitous mRNA expression, 0.5 to 1 nl of mRNA solution was injected into 1-cell stage embryos.

### Clonal analysis with Zebrabow

For clonal analysis, embryos from Zebrabow transgenic fish outcrossed to Casper fish were used. A single blastocyst per embryo was injected at the 8-16 cell stage with 6.25 pg of Cre-ERT2 mRNA and LSS-mOrange-ras mRNA. At 10 hpf, embryos were treated with 10 μM 4-OHT in E3 or 0.1% EtOH (control) for 1h, and then washed 3×5 minutes with E3. Embryos were dechorionated and fixed with 4% PFA at 24 hpf for further HuC/D immunostaining.

Confocal imaging was performed using a Zeiss780 confocal microscope with 40x/1.3 Oil DIC M27 (UV)VIS-IR Plan-Apochromat objective (FWD = 0.21 mm) with 2-μm Z intervals. Cells of the same zebrabow color (mCerulean, EYFP) that were grouped together, were spatially well-separated from other clones and labelled with the membrane-targeted were defined as a single clone. The number of neurons within a clone was determined based on HuC/D staining.

### Embryo fixation and immunostaining

Embryos were euthanized with tricaine and manually dechorionated. Embryos were fixed with 4% paraformaldehyde in PBS for 4 hours at room temperature or overnight at 4°C, and then washed 3 times for 5 minutes with PBS-TritonX 0.3% at room temperature. Embryos were permeabilzed with proteinase K and incubated with blocking solution (PBS-TritonX100 0.3% + 3% BSA) for 1 hour at room temperature. Incubation with the primary antibody (mouse monoclonal anti-HuC/D #A21271 (ThermoFisher)) was done at a concentration of 1:250 in blocking solution overnight at 4°C, followed by washing steps at room temperature. Next, embryos were incubated with secondary antibody (goat-anti-mouse Alexa647 #A21236 (ThermoFisher)) at a concentration of 1:500 in blocking solution for 2 hours at room temperature, followed by 3 5-minute washing steps at room temperature. Finally, embryos were incubated with DAPI at a concentration of 1:1000 of PBS for 48 hours at 4°C to label nuclei. After washing steps, embryos were stored at 4°C for up to one week before imaging.

### Imaging

Confocal imaging was performed at 28.5°C using an inverted Zeiss780 confocal microscope equiped with a water objective (C Apochromat"40x/1.2WCorr UVVISIR (WD=0.28mm)), at a 1-μm z-spacing along the z axis. For Cellpose data, all images were acquired at 1024×1024, except for 16 hpf stage images in dataset_1, which were taken at 512×512; at 8 or 16-bit depth with a maximum of 2-μm z-spacing.

### Image analysis

Unless otherwise specified, image processing and analysis was done using the open-source FIJI/ImageJ software (Schindelin et al., 2012). For the preparation of figures, images were adjusted for brightness and contrast, and a gaussian blur was applied (max. 1.5 pixel) using Adobe Photoshop.

### Cell counting

For the quantification of cells, confocal z-stack images showing DAPI staining were first preprocessed using the python package scikit-image (van der Walt et al., 2014). Because the image intensity is strongly reduced in the z-stack planes from the deeper parts of the tissue, contrast limited histogram equalization (CLAHE) was used to correct for this. CLAHE was done for each plane in the z-stack separately. Then, a median filter was applied to remove background noise.

The preprocessing was performed in such a way that the resulting images would have an intense, high-contrast DAPI signal throughout the z-stack, as well as overall comparable signal intensities. This was achieved by manually testing different values for the parameters defined in these functions (Suppl. Fig. 1). For CLAHE, the clip limit parameter ranged between 0.02 to 0.12, while the number of bins used was constant at 127. The footprint of the median filter was set at 5×5 pixels for all images.

The preprocessed images were then segmented using the Cellpose algorithm (Stringer et al., 2021). Segmentation was done per plane (2D) as well as in 3D (using the stitch threshold parameter). The parameters used by the Cellpose algorithm to segment the cells were adjusted to ensure a sufficient detection of the cell nuclei (Suppl. Fig. 2). The model type used was set to nuclear. The diameter parameter ranged from 25 to 30 pixels. The flow threshold ranged from 0.9 to 0.99. The cell probability threshold was set at -6. For the 3D segmentation, the automatic segmentation was turned off and the stitch threshold was set at 0.58. Implementation of the parameters was performed according to the Cellpose documentation (Stringer et al., 2021). The resulting label maps were manually checked for accuracy.

For counting the total numbers of cells in the telencephalon, image masks annotating the developing telencephalon were manually drawn using Napari (Sofroniew et al., 2022). The area in which the masks have been drawn were based on both DAPI and HuC/D staining, and the orientation of the nuclei relative to the surrounding area. The posterior borders of the telencephalon were based on the dorsal widening of the ventricle. In images where the optic recess region (ORR) emerged, the posterior borders were set such that the ORR was excluded from the telencephalon demarcation (see also Supplemental Movies 1-6). These criteria was based on publications presenting the ORR as a distinct brain region (Affaticati et al., 2015; Werner et al., 2021). Ventrally, the telencephalon borders were set by the evaginating eye vesicles. To reach a comparative relative depth of the imaging, the z-stacks were trimmed based on the widening of the ventricle at its anterior end. Images with a poor signal quality and consequently with a low accuracy of the Cellpose segmentation before the defined trim point were excluded from further analyses.

For counting neuronal numbers, the masks were manually drawn in the areas in the telencephalon with clear HuC/D labelling. These masks were first multiplied with the label maps in which the planes of the z-stack have been segmented separately. Next, the properties of each label in the resulting label map were compared with those of the originally segmented label map. All labels that had less than 80% of the size of the same label in the original label map, as well as the labels with a size of less than 300 pixels were removed. Subsequently, the resulting label map was converted into a mask and multiplied with the label map containing nuclear segmentation in 3D. The number of cells was then extracted by counting the number of unique labels present in the resulting label map.

### Division event and division mode probability calculation

For the Cellpose-based dataset, the possible total number of division events was determined based on the mean number of total cells, NPCs and neurons added within the entire telencephalon per each time interval after 16 hpf (Tables 2 and 3). Additional new NPCs are generated from symmetric proliferative division events (N_P-P_) in which one new NPC is added per division event. New neurons are born from two types of neurogenic divisions. In asymmetric division events (N_P-N_), no additional NPCs are generated but one neuron is newly generated. In symmetric terminal division events (N_N-N_), two neurons are generated at expense of one NPC.

Therefore, the number of NPCs n_NPC,*t*_ at any time *t* is:

n_NPC,*t*_ = n_NPC,*t-1*_ + N_P-P −_N_N-N_

and the number of neurons n_Neu,*t*_ at any time *t* is:

n_Neu,*t*_ = n_Neu,*t-1*_ + N_P-N +_ 2*N_N-N._

Solving these equations for time intervals 16-18, 18-20 and 20-24 hpf gives the minimum-maximum range of possible events for N_P-P_, N_P-N_ and N_N-N_ during each of these intervals.

For determining the division mode probabilities for the Zebrabow clonal analysis dataset, for each induction + chase group, the cumulative sum of possible division events leading up to that clone size and composition were determined for each individual clone according to Suppl. Table 3. For certain clone compositions when clone size ≈4, there are multiple combinations of possible division events possible to lead up to that outcome. In those cases (present only in 10-24 and 16-24 hpf datasets), the minimum-maximum range of probabilities for each division mode type was determined.

### Statistics

The coefficient of variation and coefficient of determination for the Cellpose quantification datapoints were determined using Graphpad Prism9. For comparing the total cell, NPC and neuron counts, non-parametric two-tailed Mann-Whitney test was performed using Graphpad Prism9. For comparing the Zebrabow clone type distributions between the different induction + chase groups, a 2×4 contingency table Fisher’s exact test was performed (http://vassarstats.net). P-values <0.05 were considered statistically significant. All p-values are indicated in the corresponding tables or figure legends.

## Acknowledgements

We thank Nynke Oosterhof for conceptualizing and developing the Python scripts that have been used to count the cells in the zebrafish telencephalon using the Cellpose algorithm. We thank Caren Norden and Tjakko van Ham for the kind gift of reagents and fish lines. We thank the UMCG animal facility for excellent zebrafish care. Part of the work has been performed at the UMCG Imaging and Microscopy Center (UMIC), which is sponsored by Dutch Research Council (NWO) grant 40- 175-010-2009-023.

## Author Contributions

Conceptualization: GCG, JTMLP.; Methodology: GCG, ED, CSL, YL; Experimentation: GCG, NO, ED, ESCD, HA, YL; Analysis: GCG, ED, CSL, JTMLP; Writing - original draft: GCG, JTMLP; Writing - review & editing: GCG, ED, CSL, JTMLP; Visualization: GCG, ED, CSL, JTMLP; Supervision: JTMLP; Funding acquisition: JTMLP.

## Data availability

The code will be made available on Github (github.com/nynkeoosterhof1/cell-counter.git).

