## Supplemental information for "A quantitative characterization of early neuron generation in the developing zebrafish telencephalon"

### Supplemental Figures

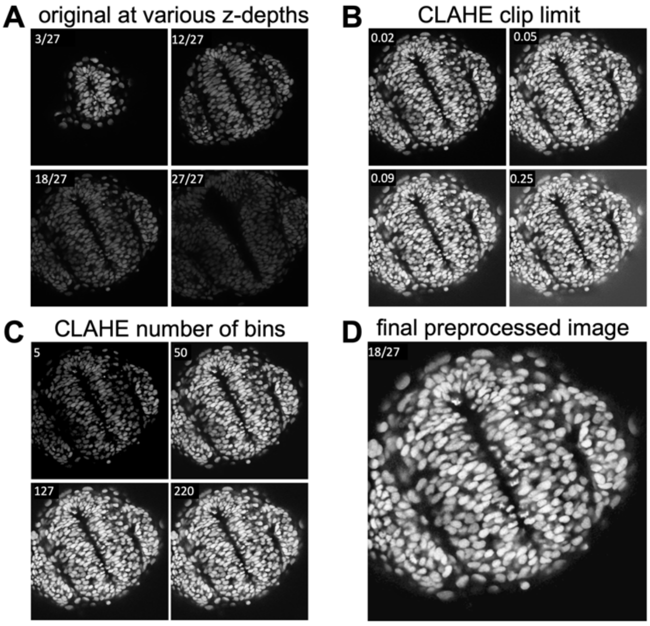


Supplemental Figure 1

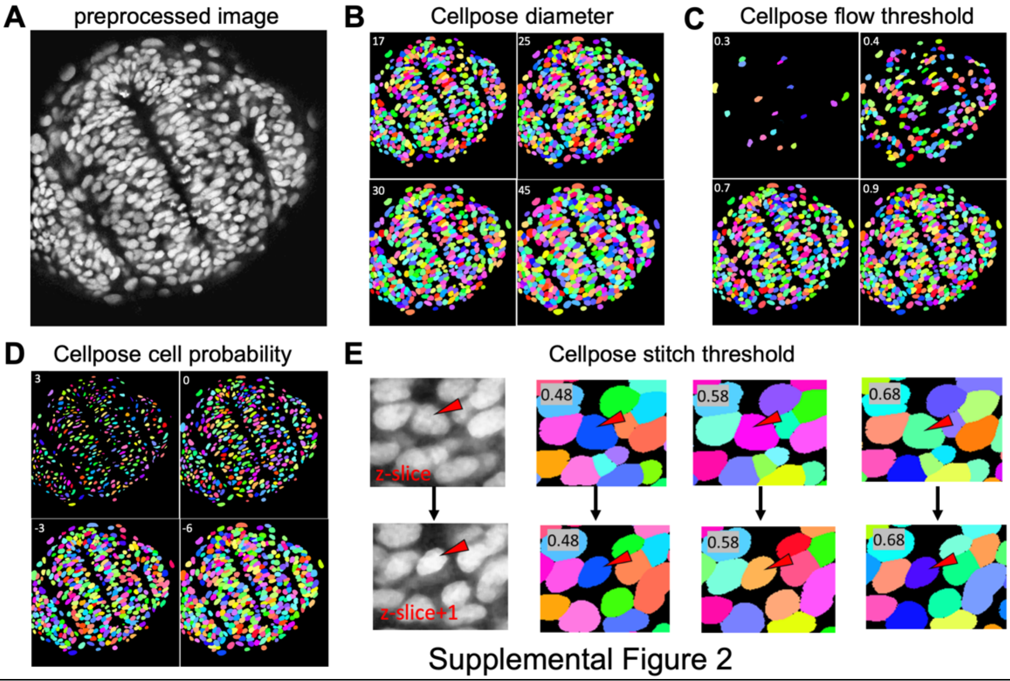


Supplemental Figure 2

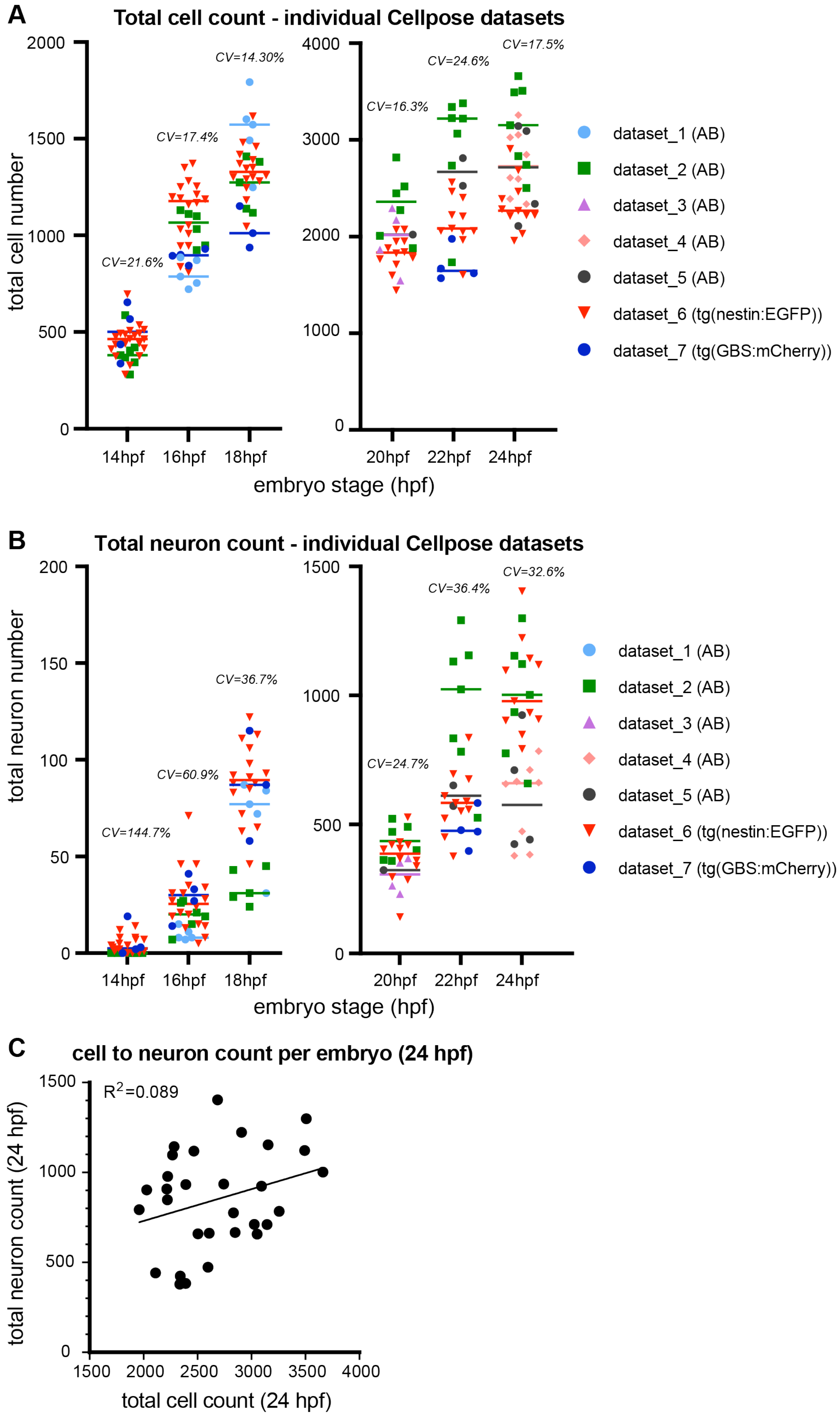


Supplemental Figure 3

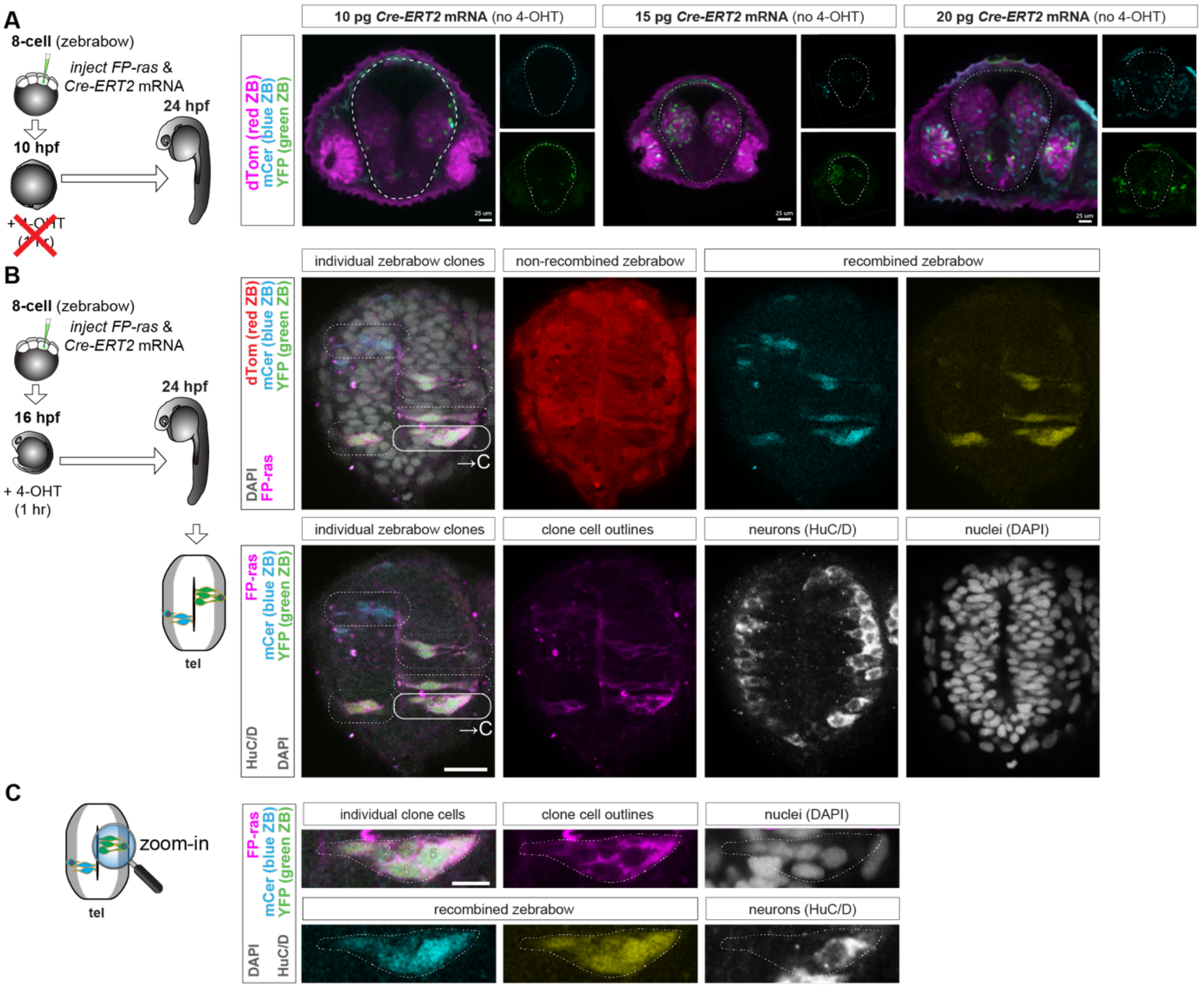


Supplemental Figure 4

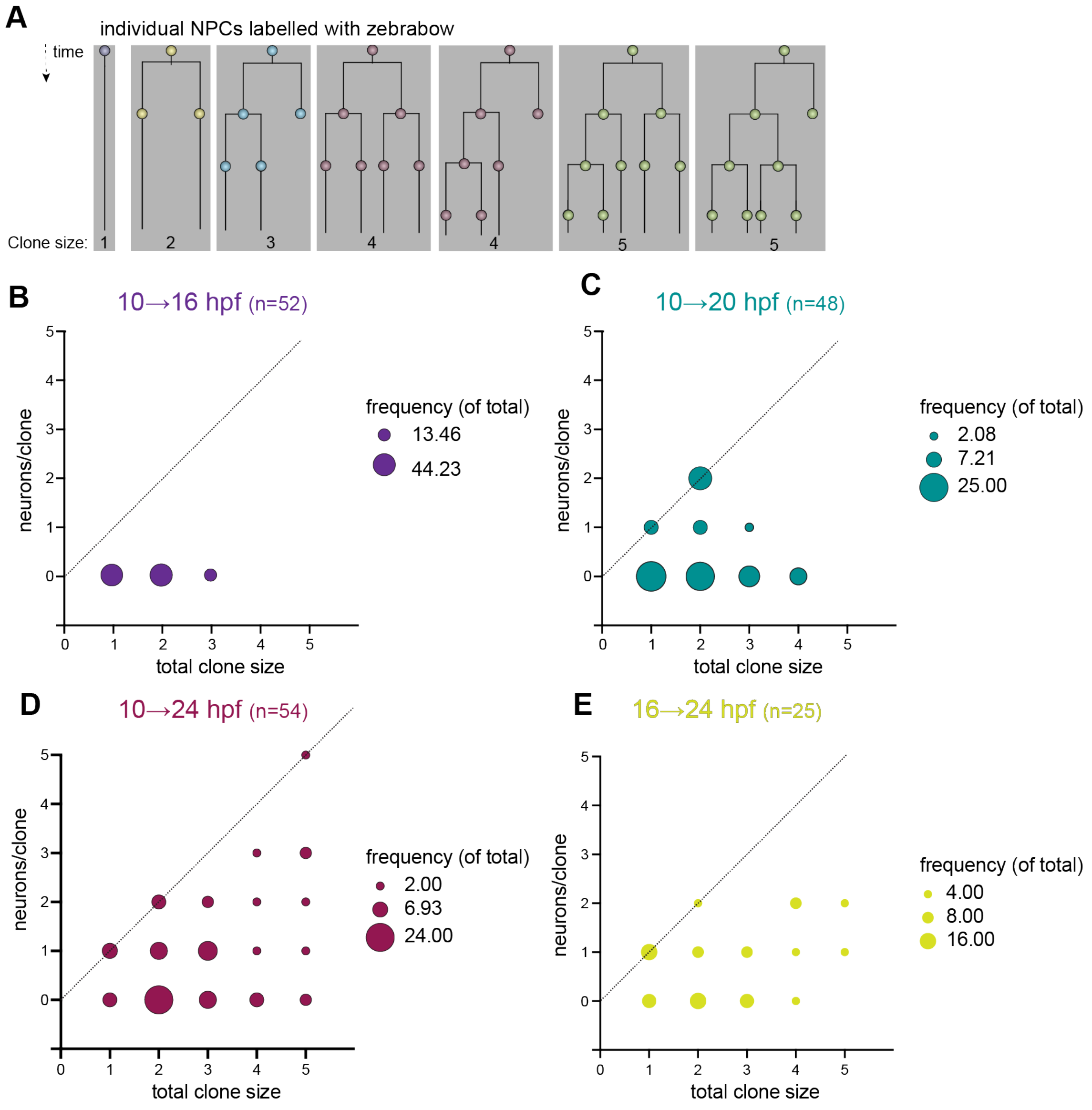


Supplemental Figure 5

**Supplemental Figure legends.**

**Supplemental Figure 1. Optimization of CLAHE parameters and their impact on the preprocessed image.** A) Exemplary original image z-slices taken from several positions throughout the z-stack. Image taken at 22 hpf. B) Side-by-side comparisons of modifying CLAHE's clip limit parameter with the number of bins set at 127 and no median filter. C) Side-by-side comparisons of modifying the number of bins used in CLAHE with the clip limit set to 0.05 and no median filter. D) Final preprocessed image slice with final settings; clip limit set to 0.05, number of bins set to 127, and median filter of 5x5 pixels.

**Supplemental Figure 2. Optimization of Cellpose parameters and their impact on the nuclear segmentation.** A) Exemplary slice of a preprocessed image to run through the Cellpose nuclear segmentation algorithm. Image taken at 22 hpf. B) Comparison of various diameter values at a flow threshold of 0.9 and a cell probability threshold of -6. C) Comparison of various flow threshold values at a diameter of 25 and a cell probability threshold of -6. D) Comparison of various cell probability threshold values at a diameter of 25 and a flow threshold of 0.9. E) Selection of two consecutive slices within the z-stack. Arrows shows two nearby cells along the z-stack. Comparison of various stitch threshold values shows that the cells are mislabeled as one cell for a stitch threshold of 0.48, while other parameter values correctly identify them as two distinct cells. The same color indicates that the cell is segmented as one entity by Cellpose.

**Supplemental Figure 3. Quantification of cell numbers for each individual dataset.** Cellpose-based quantification of the total number of cells A) and B) neurons present in the telencephalon for 14 to 18 hpf (left) and 20 to 24 hpf (right) as determined using the Cellpose pipeline. Note that the y-axis is rescaled between left and right graphs to facilitate interpretation. Each point reflects data from one individual embryo; the individual datapoints from each individual dataset are marked with different color and marker shapes according to the index. The colored lines indicate the median for each individual dataset. The coefficient of variation (CV) is calculated based on all individual datapoints combined. C) The total number of cells and neurons in the telencephalon of individual embryos at 24 hpf. A linear regression model goodness-of-fit was performed with the coefficient of determination ($R^2$) indicated.

**Supplemental Figure 4. Clonal analysis of NPCs in the developing telencephalon using zebrabow recombination.** A) Representative (single-optical section) confocal image of the frontal view (non-recombined zebrabow cassette (dTom, magenta), recombined zebrabow cassette mCer, cyan; EYFP; green)) of the telencephalon (outlined with dashed line) of *ubi*:zebrabow embryos in which different

*CreERT2* mRNA amounts (10-20 pg) were injected to find mRNA concentration with no to low recombination in the absence of 4-OHT treatment. Dorsal is up. Amount of injected *CreERT2* mRNA is indicated. Insets show single fluorescent channels for recombined cassettes (mCer cyan; EYFP; green). B) Representative (single-optical section) confocal image of the frontal view of the telencephalon of a 24 hpf zebrabow embryo, induced at 16 hpf, showing several individual clones (indicated with dashed lines) for individual and overlay fluorescent signal channels for non-recombined zebrabow (dTom, red), recombined zebrabow cassette mCer, cyan; EYFP; yellow), membrane-targeted LSSmOrange-ras (magenta), neurons (grey) and nuclei (grey). Dorsal is up. Scale bar 25μm. One of the clones (white line) is magnified in C). C) Magnified view of a five-cell clone, showing the overlap of one of the cells with HuC/D. Scale bar 10 μm.

**Supplemental Figure 5. Clonal composition of zebrabow induced and chased groups.** A) Theoretical individual NPC lineage trees that lead to each observed clone size. Each branch indicates a cell division event. B-E) Overview of the clonal composition (neurons/clone) per clone size expressed as frequency (percentage of all clones) for each 10 to 16 (B), 10 to 20 (C), 10 to 24 (D) and 16 to 24 (E) hpf induction + chase groups. The size of each datapoint indicates the frequency of that particular clone composition/clone size within all the clones.

**Supplemental Movies 1-6.**

Example images of cell nuclei (left) and neuronal (right) segmentation by Cellpose within the telencephalon as determined by manually drawn masks for the telencephalon region. Shown are segmented nuclei outlines (white lines) overlaying DAPI (grey) signal (left) and neuronal outlines overlaying HuC/D immunofluorescence (grey) signal (right) for 14 hpf (Movie 1), 16 hpf (Movie 2), 18 hpf (Movie 3), 20 hpf (Movie 4), 22 hpf (Movie 5) and 24 hpf (Movie 6).

**Supplemental Table 1. Raw data Cellpose-based quantifications (xls file)**
**Supplemental Table 2. Raw data Zebrabow clones (xls file)**
**Supplemental Table 3. Potential division events in the lineage for each clone type/size.**

| Clone size | Clone composition (# of neurons) | $N_{P-P}$ (symmetric P-P division) | $N_{P-N}$ (P-N division) | $N_{N-N}$ (symmetric N-N division) |
|---|---|---|---|---|
| 1 | 0 | 0 | 0 | 0 |
| 1 | 1 | 0 | 0 | 0 |
| 2 | 0 | 1 | 0 | 0 |
| 2 | 1 | 0 | 1 | 0 |
| 2 | 2 | 0 | 0 | 1 |
| 3 | 0 | 2 | 0 | 0 |
| 3 | 1 | 1 | 1 | 0 |
| 3 | 2 | 0 | 2 | 0 |
| 3 | 3 | 0 | 1 | 1 |
| 4 | 0 | 3 | 0 | 0 |
| 4 | 1 | 2 | 1 | 0 |
| 4 | 2 | 1 or 2 | 0 or 2 | 0 or 1 |
| 4 | 3 | 1 | 1 | 1 |
| 4 | 4 | 0 or 1 | 0 or 2 | 1 or 2 |
| 5 | 0 | 4 | 0 | 0 |
| 5 | 1 | 3 | 1 | 0 |
| 5 | 2 | 1 or 2 or 3 | 0 or 2 | 0 or 1 |
| 5 | 3 | 1 or 2 | 1 or 2 or 3 | 0 or 1 |
| 5 | 4 | 1 | 1 or 2 | 1 or 2 |
| 5 | 5 | 1 | 1 | 2 |
